# Speech Stream Tracking in 2D: Attention Differentially Enhances Acoustic and Phonemic Encoding Across Spatial Planes

**DOI:** 10.64898/2026.06.03.729740

**Authors:** Varvara Kenti Kranidioti, Marc Schönwiesner

**Affiliations:** Institute of Biology, Faculty of Life Sciences, University of Leipzig, Leipzig, Germany; International Laboratory for Brain, Music, and Sound Research, Montreal, Quebec, Canada

**Author notes:** **Correspondence:** Marc Schönwiesner.

**Keywords:** EEG, selective attention, temporal response function, stream segregation, spatial hearing, phoneme encoding

## Abstract

Selective attention enables the flexible allocation of neural resources toward relevant stimuli. In audition, it allows listeners to track a target stream within acoustically complex environments. This raises the question of how strongly auditory attention depends on spatial cues to achieve stream segregation and maintain distinct auditory objects. Evidence shows that attention enhances neural encoding of sound features for streams separated in azimuth. However, it remains unclear whether the same mechanisms apply without binaural cues in elevation, and how attention prioritizes acoustic versus linguistic features under these conditions. To address this, participants listened to arrhythmic streams of digits spoken by the same voice and separated in azimuth or elevation. They attended one stream to detect target numbers while ignoring the other. Neural responses to attended and ignored streams were modelled using envelope and phoneme temporal response functions, allowing comparison of low-level envelope and higher-level phonemic encoding across spatial dimensions. Results revealed distinct feature-weighting profiles across spatial planes. In azimuth, selective attention was primarily supported by enhanced envelope encoding of the target stream at early and middle latencies. In elevation, envelope encoding was reduced, while phoneme encoding exhibited a more widespread attentional modulation. These findings suggest that phonemic representations support selective stream tracking when binaural spatial cues are unavailable, reflecting flexible weighting of acoustic and phonemic information depending on spatial cue availability.

## INTRODUCTION

Our world is filled with an abundance of sensory input, yet despite a constant and sometimes overwhelming amount of information, we manage to focus on a selected stream while ‘ignoring’ the rest. This ability is called selective attention: the allocation of neural resources to relevant stimuli. Although attention functions as a global mechanism across sensory systems, its influence on neural activity reflects an interplay between supramodal control processes and system-specific sensory modulation (Macaluso et al., 2002; Shomstein & Yantis, 2004; Fu et al., 2020).

Unlike vision, where directing gaze provides inherent spatial enhancement (Daniel & Whitteridge, 1961; Dow et al., 1981), auditory attention must operate on streams that overlap at the sensory epithelium, a challenge described as the “cocktail party problem” (Cherry, 1953). Although stream segregation in complex auditory scenes can begin preattentively (Bregman, 1990; Sussman et al., 1999; Sussman et al., 2005), sustained selection of a target stream depends on attention, which enhances and refines auditory object representations over time (Best et al., 2008; O’Sullivan et al., 2015). Evidence shows that selective attention enhances neural encoding of both low-level acoustic features, such as the speech envelope (Ding & Simon, 2012; Golumbic et al., 2013), and higher-level linguistic features, including phonemic and semantic structure (Di Liberto et al., 2015; Broderick et al., 2018).

Most studies on selective attention to competing speech streams have focused on sources arranged along azimuth (Ding & Simon, 2012; Power et al., 2012; Golumbic et al., 2013; O’Sullivan et al., 2015; Di Liberto et al., 2015; Broderick et al., 2018; Kaufman & Golumbic, 2023). There, robust binaural cues support sound localization and stream segregation (Middlebrooks & Green, 1991; Blauert, 1997; Deng et al., 2019). Under optimal conditions, listeners can discriminate interaural time differences (ITDs) as small as ∼10 μs and interaural level differences (ILDs) of ∼0.5 dB (Mills, 1958), making binaural cues highly effective for separating simultaneous streams. These cues are extracted in the superior olivary complex in the lower brainstem (Masterton & Diamond, 1967; Yin & Chan, 1990), contributing to partially separable neural representations of competing spatial streams already in primary auditory cortex (Yao et al., 2015). By contrast, sounds originating from different elevations along the midline are discriminated primarily by monaural spectral-shape cues imposed by the pinnae (Batteau, 1967; Wightman and Kistler, 1989), likely processed in secondary or higher auditory cortex areas (Xu et al., 1998). Despite its ecological importance, selective attention to vertically separated speech is virtually unexplored.

In azimuth, an attended speech stream can be tracked with EEG based on the sound envelope, suggesting that the brain segregates competing streams using low-level acoustic cues, potentially supported by early, non-overlapping neural representations. However, such separable representations of acoustic features are likely weaker for streams separated in elevation. We therefore hypothesize that speech stream segregation in elevation relies stronger on phonemic than acoustic features. This distinction between binaural and spectral spatial cues provides a framework to test whether the auditory system differentially prioritizes envelope versus phonemic information, depending on spatial cue availability, revealing how feature weighting supports selective tracking.

We compared neural and behavioral responses to competing speech streams separated in azimuth or elevation. Both streams were produced by the same talker and presented with arrhythmic timing, eliminating voice-specific cues and temporal predictability as segregation features. We quantified encoding of envelopes and phoneme onsets for target and distractor streams using temporal response function (TRF) modelling. Neural coupling strength was quantified using a correlation index (Δr), defined as the difference in prediction accuracy between target and distractor streams, and related to behavioral performance using mixed-effects regression. We tested three hypotheses: (1) envelope target–distractor encoding is stronger in azimuth than elevation, reflecting robust binaural segregation; (2) phoneme-based selectivity is relatively stronger in elevation, reflecting increased reliance on linguistic structure when spatial cues are weaker; and (3) neural selectivity predicts behavioral performance, indicating functional relevance for stream segregation.

## MATERIALS AND METHODS

### Participants

Twenty-one healthy participants (15 women, 6 men; age range: 18–40 years, mean age: 28.4 years, SD = 5.07 years) took part in the study. Sample size was determined based on prior TRF studies (e.g., Ding & Simon, 2012, *N* = 11; Di Liberto et al., 2015, *N=*10) and is consistent with recommended sample sizes for EEG TRF paradigms (Crosse et al., 2016, *N* = 18). All subjects reported normal hearing. Participants received monetary compensation for their time. Before the experiment, all participants read the study’s procedure and details and provided written consent. Any remaining questions were addressed, and participants underwent a brief training session during which they listened to a short experimental block and practiced responding to the target stream to ensure full understanding of the task. The procedure was approved by the Ethics Committee of the University of Leipzig. Three participants were excluded from further analysis due to issues affecting data validity. One participant was excluded due to behavioral performance indicating failure to comply with task instructions. Two participants were excluded due to poor EEG signal quality caused by a recording hardware malfunction. The final sample consisted of eighteen participants included in the analysis.

### Stimuli and Speech Stream Design

High-quality neural text-to-speech (TTS) recordings (two male, two female voices) were used. The selected voices were perceptually natural, with clear prosody and no synthetic artifacts or background noise. The stimuli consisted of the number words 1 through 9, excluding the two-syllabic 7.

Each stream consisted of pseudorandomized sequences of spoken numbers. Simultaneous presentations of the same number across both streams were avoided, ensuring that responses to the distractor are not mistakenly classified as target responses, and vice versa. All numbers appeared with equal probability (12.5% of total trials) in both streams. A target number was randomly selected in each block and presented more often (35% of total trials) in the target stream. Additionally, animal sounds were played in the distractor stream (10% of total trials) as deviant stimuli to enhance the stream’s effectiveness. For this, nine isolated animal vocalizations were used, extracted from public-domain video sources. The sounds were cleaned and edited in Audacity, where background noise was attenuated. These events did not replace target-number occurrences, and they were discarded prior any analysis. All stimuli were 745 ms long, sampled at 24.414 kHz, and played at 80 dB SPL.

Target and distractor streams were presented at different inter-stimulus intervals (ISI) of 90 ms or 70 ms. Additionally, a variable delay, randomly drawn on each trial from a uniform distribution between 0 and 10 ms, was added to the ISI. This design prevented participants from entraining themselves to the rhythm and encouraged active monitoring of the target stream. To indicate the target stream to the listener, the distractor stream was delayed by 3 stimulus presentations (approx. 2.5 s) relative to the target stream. Each block lasted 2 minutes.

Each participant completed 20 blocks in total, divided evenly across four spatial attention conditions (five blocks per condition, ∼10 min. each). The number of stimuli in each condition’s stream was consistent across participants and spatial planes (709.5±6.45 stimuli, <1% variability). The order of spatial conditions was randomized across participants, and voices were counterbalanced to avoid systematic bias. In the azimuth conditions, the target and distractor streams were presented from lateral loudspeakers. Depending on the block, participants were instructed to attend either to the right or to the left stream while ignoring the other. In the elevation conditions, the two streams were presented along the vertical midline, and participants were instructed to attend either the upper or the lower stream accordingly. Attended location was counterbalanced across blocks to ensure unbiased spatial cueing.

### Experimental Procedure

Participants were seated in a hemi-anechoic chamber facing the loudspeakers, with their ear-level aligned with the midline loudspeaker, 1.2 m above ground. Communication with the experimenter was possible via an intercom system in the control room.

Prior to block initiation, the target number was announced, and verbal confirmation was obtained by the participant. Then the experiment began with the target stream, followed by the distractor with a ∼2.5 s delay, giving the participants enough time to focus on the target stream, before both streams played concurrently. No target number was presented in the target stream before the onset of the distractor stream. During each block, participants selectively attended to the target stream while ignoring the distractor, pressing the corresponding key on a number pad as quickly as possible whenever they heard the target number. All stimulus onsets and button presses were time-locked and stored as event markers within the EEG recording. Trial sequences were generated using the slab Python module (Schönwiesner and Bialas, 2021).

### EEG Acquisition

Subjects sat in the center of a custom-built spherical array of 48 loudspeakers (model Mod1, Sherman Oaks, CA, USA) inside a 40 m2 hemi-anechoic chamber (Industrial Acoustics Company, Niederkrüchten, Germany). Loudspeakers were equidistant (1.4 m) from the participant’s head. For this experiment, only four speakers were used (at ±6.5° azimuth and ±37.5° elevation). The angular separations were to account for the better discrimination threshold in azimuth even at small separations, whereas elevation required wider spacing to achieve reliable performance. An acoustically transparent curtain concealed the speakers from the participants. We equalized the transfer functions of each loudspeaker by applying an inverse filter to the stimuli upon presentation. Two digital signal processors and six 8-channel amplifiers (models RX8 and SA8, Tucker-Davis Technologies, Alachua, FL, USA) drove the loudspeakers at 24.414 kHz sampling rate. The processors’ digital ports obtained responses from a custom-built button box and sent event triggers to the EEG. We used a 64-channel system (model BrainAmpDC, Brain Products, Gilching Germany) to record EEG at a sampling rate of 500 Hz. The active silver/silver chloride electrodes were fixed on the subject’s head with an elastic cap (Easycap, Germany) according to the international 10/20 system with FCz as the online reference electrode. Electrode impedance was kept below 10 kΩ.

### EEG Pre-processing

EEG data were preprocessed in Python using the MNE toolbox (Gramfort et al., 2013). A zero-phase finite-impulse-response band-pass filter (1-30 Hz) was first applied to remove slow drifts and high-frequency noise. The filtered data were visually inspected and segments showing global artifacts or large non-physiological fluctuations (e.g., due to muscle or reference noise) were manually marked for exclusion. Channels exhibiting persistent noise or dropout were also marked for later interpolation. Independent Component Analysis (ICA) was then performed on the artifact-masked data to identify and remove components corresponding to ocular activity, including eye blinks and saccades. Only oculomotor components were removed. The cleaned data were subsequently low-pass filtered at 8 Hz, following Di Liberto et al. (2015), who demonstrated that low-frequency cortical activity carries robust speech-related information. The data were then re-referenced to the common average and downsampled to 125 Hz.

### TRF Modeling

EEG responses were modeled from time-resolved stimulus features using a forward encoding temporal response function (TRF; Bialas et al., 2023) approach. For each participant, spatial condition (azimuth or elevation), and stream (target or distractor), a linear mapping between time-resolved stimulus features and the corresponding EEG signal were estimated. Three types of predictors were included. (1) Speech envelopes: For each stream, amplitude envelopes were extracted following a short-term root-mean-square procedure adapted from Power and colleagues (2012). Root-mean-square energy was computed in 195-sample windows (matching the EEG sampling interval of 8 ms), smoothed over a 390-sample window, and resampled to 125 Hz. (2) Phoneme onsets: The spoken digits were aligned using the Montreal Forced Aligner (McAuliffe et al., 2017). For each token, phoneme onset times were extracted and converted into sparse binary impulse trains (1 = onset, 0 = otherwise), aligned with the EEG data. The first phoneme of each word, which is highly correlated with envelope transients, was excluded, to avoid confounding the model with low-level acoustic onsets, ensuring that phoneme predictors captured within-word linguistic structure. Separate phoneme regressors were constructed for the target and distractor streams. (3) Button presses: Behavioral responses were encoded as binary impulses aligned with each key press and included as a nuisance regressor to account for motor-related EEG variance.

Artifact-marked EEG segments (< 1% of total data) were masked in both EEG and predictor time series to maintain temporal alignment. Finally, the data were z-scored per channel.

Separate TRF models were computed for azimuth and elevation blocks. For each participant, a design matrix was constructed by concatenating predictors from target and distractor streams column-wise, allowing the model to simultaneously estimate neural responses to both streams, while accounting for shared variance between predictors. Envelope predictors were z-scored. When only the very sparse target-number onsets were modeled, instead of z-scoring, the non-zero elements were mean-centered. The resulting predictor matrix (X; time × predictors) and EEG data (Y; time × channels) were entered into a TRF model estimated with ridge L2 regularization (mTRFpy Toolbox for Python; Bialas et al., 2023). The lag window spanned -100 to +1000 ms relative to each predictor time point, encompassing a 100 ms pre-stimulus interval, and an extended post-stimulus period to capture late auditory and cognitive responses. The extended lag window reduces temporal truncation of overlapping neural responses in continuous stimulus streams with short inter-stimulus intervals. All statistical analyses were restricted to post-stimulus activity in a 0 to 500 ms latency window.

A single global regularization parameter was selected to ensure comparability across participants, conditions and planes and to prevent overfitting. For each condition, predictor matrices (X) and EEG data (Y) were assembled for each participant. Regularization strength was optimized using ridge L2 regression across a logarithmically spaced range of λ values (0.01–100). For each λ, prediction accuracy was estimated using 5-fold cross-validation across contiguous, non-overlapping temporal segments of the continuous data. In each fold, models were trained on 80% of the data and evaluated on the remaining 20%. Prediction accuracy on held-out data was quantified as the Pearson correlation (r) between predicted and recorded EEG signals, computed per channel and averaged across folds. Cross-validated accuracy values were then averaged across channels and participants to obtain a global performance estimate for each λ. Prediction accuracy reached its maximum at λ = 0.01 and remained stable across cross-validation folds, indicating that this value provided sufficient regularization while preserving generalization performance. Therefore, λ = 0.01 was used all subsequent TRF model fits. Final model prediction accuracy was then computed with the fixed λ and the same 5-fold cross-validation procedure for each participant, computed independently for each EEG channel and averaged within the predefined ROI, providing a subject-level measure of neural encoding strength. All TRF weights and subject-wise prediction accuracies were retained for group-level statistical analyses and visualization.

To confirm that prediction accuracy reflected genuine stimulus-brain coupling, control (“null”) TRF models were trained using circularly time-shifted predictors, providing a conservative baseline while preserving predictor autocorrelation structure. For each subject, every continuous predictor (speech envelopes, phoneme onsets, and button-press impulses) was circularly shifted by a random lag between 0.5 and 5 s, independently for each predictor and stream. This procedure preserved each predictor’s autocorrelation structure but disrupted both intra- and inter-stream temporal alignment with the EEG. Null models were fit using the same λ and lag window as the real models.

### Statistical Analysis

Regions of interest (ROIs) were selected based on previous literature. For the analysis of envelope TRF encoding, the central channel Cz was used, consistent with prior studies demonstrating robust speech envelope-following responses over frontocentral and central midline scalp regions, including Cz (Aiken & Picton, 2008; Lalor & Foxe, 2010; Ding & Simon, 2012). For phonemes, a 12-channel frontotemporal ROI (F3–F8, FC3–FC6, FT7–FT8) was predefined, aligned with prior phoneme-tracking work showing maximal phoneme-level responses (Di Liberto et al., 2015). Complementary whole-head cluster permutation tests did not identify stable channel clusters, supporting the use of the literature-driven ROI for primary statistical analyses. Additionally, whole-scalp prediction accuracy was broadly distributed across electrodes (target model: M = 0.189, SD = 0.031; distractor model: M = 0.180, SD = 0.030. SD reflects variability across electrodes), with low and consistent inter-subject variability (mean SD across subjects ≈ 0.05). This indicates that stimulus encoding was not restricted to a small subset of channels, supporting the use of literature-based ROIs rather than selecting electrodes based on consistently high prediction accuracy across conditions and participants. To assess whether the predefined ROIs adequately captured the spatial distribution of encoding effects in the dataset, complementary robustness analyses were conducted using modestly expanded ROIs. These exploratory ROIs were defined to reflect the spatial distribution observed in grand-average scalp topographies (see supplementary Figures S1 and S2) and to test whether conclusions were robust to modest ROI changes, and not to optimize statistical significance.

TRF phoneme encoding was standardized to account for the binary and sparse nature of the predictor. Each subject’s TRF weights were multiplied by *sqrt(p(1−p))*, where *p* is the proportion of non-zero impulses in the predictor, yielding coefficients scaled to the predictor variance. Because EEG data were z-scored, the resulting β values reflect the standardized EEG change per unit change in predictor activity.

Four different latency windows were defined a priori (50–150 ms, 150–250 ms, 250–350 ms, and 350–500 ms) to reflect successive stages of auditory processing from early sensory responses to later cognitive or linguistic evaluation. Testing within latency ranges reduces the likelihood that broad clusters spanning most of the epoch obscure stage-specific effects and allows targeted interpretation of early versus late attentional modulation. Whole-epoch analyses were additionally performed to confirm that effects detected in latency-window analyses remained detectable under conservative correction across the entire temporal search space.

Statistical comparisons of ROI-averaged TRF weights were performed between target and distractor streams within each condition, using non-parametric cluster-based permutation tests implemented in the MNE-Python toolbox (Gramfort et al., 2013). Tests were two-tailed with 1000 permutations and a cluster-forming threshold of p < .05, applied across timepoints within each latency window. Cluster significance was determined using a maximum cluster-mass statistic, which controls the family-wise error rate across timepoints. The resulting *p*-values were corrected for multiple comparisons across components using the false discovery rate (FDR; Benjamini–Hochberg, 1995). Per subject, target–distractor amplitude differences were computed for each temporally significant cluster and visualized as topographical inset maps within the TRF amplitude plots.

Permutation tests were also used to assess whether the magnitude of target–distractor separation differed between azimuth and elevation. For each subject, difference waves between target and distractor streams were computed separately for envelope and phoneme TRF weights in the azimuth and elevation conditions. The subject-wise difference waves were compared across planes using cluster-based permutation tests within each predefined latency window.

### Brain-behavior relationships

For the subsequent brain-behavior correlation analysis, we computed performance accuracy as the behavioral measure and the difference in TRF encoding weights between streams as the neural measure.

Performance accuracy was estimated from the participants’ button responses. Those occurring within 0.2–0.9 s after target onset were classified as valid, all others as invalid. Hit rate (HR) was computed as *number of valid responses / total number of target trials*. Miss rate (MR) was computed as *number of missed targets / total number of target trials*. Responses to the target digit occurring in the distractor stream were considered false alarms and a false alarm rate (FAR) was computed as *number of false alarms / total number of distractor numbers trials*. A raw performance score was then computed as HR – MR – FAR. This composite performance metric preserves graded behavioral variability and directly reflects attentional performance, by jointly accounting for correct detections, misses, and false alarms. This provides a sensitive measure of behavioral variability suitable for regression analyses relating neural and behavioral measures. To have behavioral and neural measures on a comparable scale, performance scores were z-scored across participants within each condition.

To quantify attentional enhancement in TRF encoding the target and distractor correlation values (r, Pearson correlation between predicted and actual EEG for each stream) were obtained from reduced TRF models, each containing only one stream’s predictors, allowing independent estimation of target and distractor encoding strength without shared variance between competing predictors. Predictor collinearity was assessed using variance inflation factors (VIF), which were low across predictors, indicating minimal multicollinearity (VIFs ≈ 1.00). To ensure an approximately normal distribution, the r values were transformed using Fisher’s formula, *r*_*z*_ = tanh^-1^ *r*. The average difference between transformed target and distractor correlations (Δ*r* = *r*_*z,target*_ − *r*_*z,distr*_) was extracted from the designated ROI for each predictor type (phonemes and envelopes) and analyzed in separate mixed-effects models for the azimuth and elevation conditions. Linear mixed-effects models were implemented in Python using the statsmodels package. The following model was used for each plane:

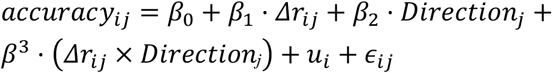

where *i* and *j* index subjects and within-plane sub-conditions respectively. Δr is entered as a fixed effect, and the categorical term *Direction* codes the two spatial sub-conditions within each plane: target-left vs. target-right for azimuth, and target-bottom vs. target-top for elevation. The interaction term assesses whether the strength of the Δr-performance accuracy relationship differs between these within-plane directional configurations. A random intercept *u*_+_ was included for each participant to model inter-individual differences in baseline performance. Random slopes were evaluated but excluded due to limited observations per subject and no improvement in model fit. All regressors were mean-centered within each plane to facilitate interpretation and reduce collinearity. Model coefficients (β) were evaluated using Wald *z*-tests with associated standard errors, 95% confidence intervals, and *p*-values. Significance was set at *alpha* < 0.05. Both models were estimated using maximum likelihood (ML). Prior to the final LMM analysis, the model’s residuals were examined to detect any significant outliers. One outlier was identified in the azimuth dataset, showing deviations from normality (Shapiro–Wilk p = 0.004). After exclusion (|z| > 3 SD), residual distributions for both azimuth and elevation models met normality and homogeneity of variance assumptions (all *p* > 0.09). All p-values from the LMM results were corrected for multiple comparisons with FDR.

## RESULTS

Participants performed highly accurately overall, with reliable task engagement across both spatial planes. In azimuth, performance reached near-ceiling levels: the mean hit-rate was 97.28% and false alarms were very low (1.54%). Elevation trials were more demanding, as expected from classic spatial-acuity asymmetries, yet accuracy remained high for most participants. The mean hit-rate was 89.60%, with half of the sample performing above 90% and only one participant falling below 80%. False-alarm rates showed more variability but were still low for the majority: 11 out of 18 listeners had values below 10%, and only one participant showed an unusually high false-alarm tendency. This pattern suggests that participants were able to carry out the task well in both planes, and that the azimuth–elevation difference arose primarily from increased distractor responses rather than failures in target detection.

### Real TRF Models Outperform Null Models

Prediction accuracy was higher for real TRF models than for circularly time-shifted controls across predictor types and spatial planes, confirming that the models captured genuine stimulus–brain coupling. In the azimuth plane, phoneme models showed significantly higher prediction accuracy than null models (real M = 0.184, null M = 0.102), W = 0, p < .001, effect size r_e_ = 1.00. Envelope models showed an even larger difference (real M = 0.223, null M = 0.116), t(17) = 8.52, p < .001, d = 2.01. A similar pattern was observed in elevation. Phoneme models showed higher prediction accuracy than null models (real M = 0.179, null M = 0.101), t(17) = 15.84, p < .001, d = 3.73, and envelope models also showed significantly higher prediction accuracy than null models (real M = 0.215, null M = 0.119), W = 0, p < .001, effect size r_e_ = 1.00. Additionally, inter-subject variability of real-model prediction accuracy was lower for phoneme models than for envelope TRF models in both spatial planes (phonemes: SD = 0.035– 0.041; envelopes: SD = 0.062–0.066), indicating that any reduced phoneme selectivity effects in elevation are unlikely to reflect inter-individual variability alone.

### Spatially Distinct Temporal Clusters in Envelope and Phoneme TRF Responses

In azimuth, cluster-based permutation tests revealed that TRF envelope encoding differed between target and distractor streams (Fig. 1A) in an early (60–91 ms, p_FDR_ = .022, g_z_ = −1.05), middle (150–200 ms, p_FDR_ = .001, g_z_ = 1.17), and late time window (260–320 ms, p_FDR_ = .001, g_z_ = −1.11). The middle cluster also emerged in a further comparison of non-target and target-number stimuli within the attended stream. In elevation, significant differences emerged only in the late time window from 300 to 340 ms (p_FDR_ = .006, g_z_ = −0.90; Fig. 1B).

**Figure 1:**
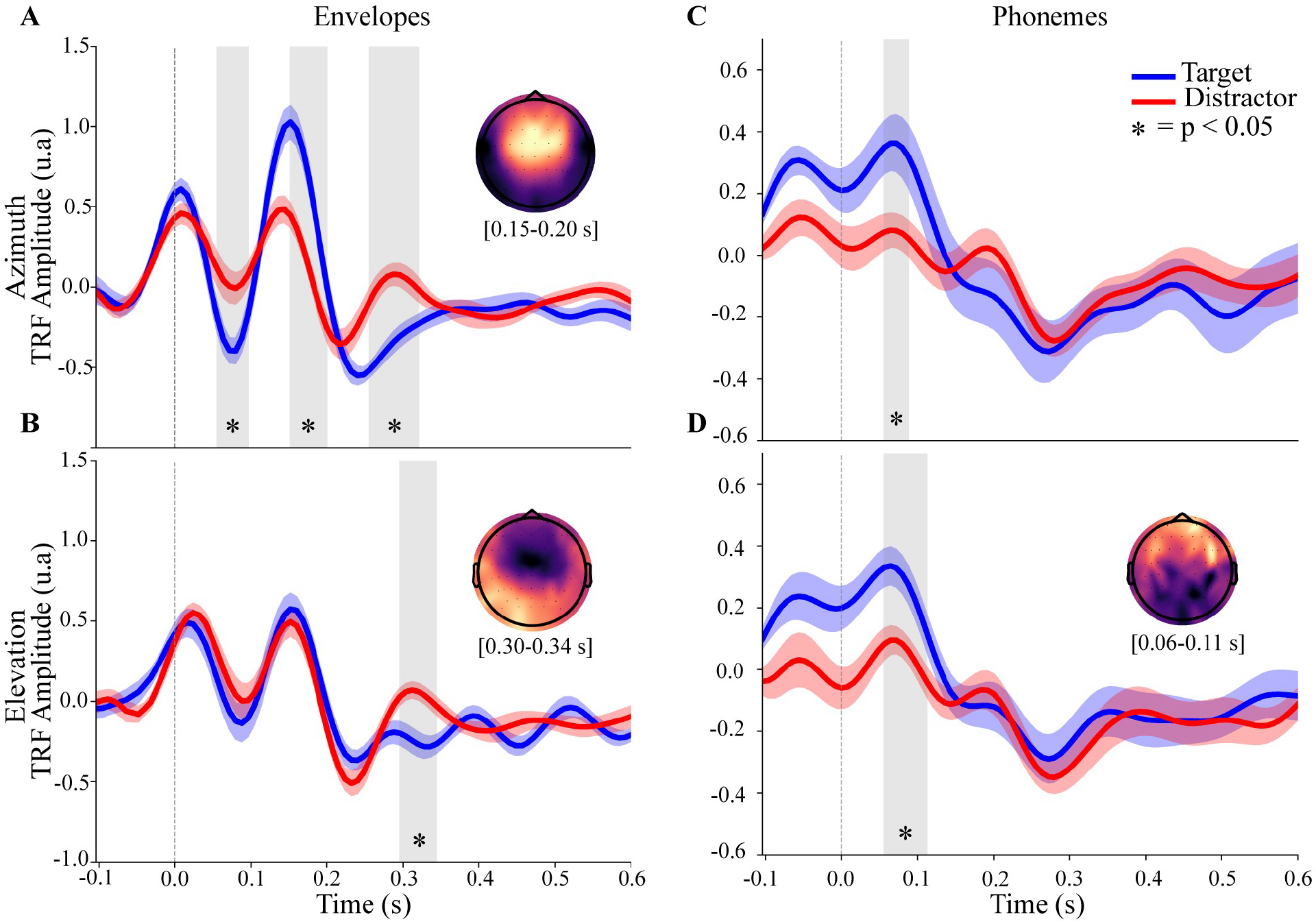
Temporal differences in TRF envelope and phoneme encoding across attentional streams. Panels A-D show the reconstructed TRF amplitudes to envelopes **(A-B)** and phoneme onsets **(C-D)** for target (blue) and distractor (red) streams in the azimuth (top row) and elevation (bottom row) conditions. For these reconstructions, all stimulus events associated with each stream were included (target and non-target stimuli). The y-axis reflects response magnitude (arbitrary units; based on standardized EEG and predictor data), and the x-axis denotes time relative to stimulus onset (-0.1 to 0.6 s). Significant differences between target and distractor responses, identified via cluster-based non-parametric permutation testing, are highlighted by shaded gray time windows (*p* < .05, FDR-corrected). Insets show scalp topographies of these significant clusters (target - distractor), averaged across participants, with red indicating target > distractor and dark purple indicating distractor > target; brackets beneath each topomap indicate the corresponding latency range, rounded to two decimal places. In Panel A, the embedded topography reflects the average amplitude differences between target and distractor responses within the 0.15-0.20 s time-window. The comparisons were conducted using the main, frontotemporal phoneme ROI (F3–F8, FC3–FC6, and FT7–FT8). Envelope analyses were performed using a Cz ROI.

The TRF encoding to phoneme onsets also differed between streams in both azimuth and elevation, with significant clusters emerging in early latencies (all p_FDR_ < .05; Fig. 1C–D). Breaking the responses down by stimulus type (task-relevant targets vs. task-irrelevant non-targets) revealed distinct differences between stimulus categories and spatial planes (see Fig. 2). In azimuth, the effect was primarily driven by target stimuli (p_FDR_ = .002, g_z_ = 1.00; Fig. 2A). In elevation, each stimulus type produced a distinct pattern of significant clusters across time. Task-irrelevant stimuli showed an early-latency cluster (g_z_ = 0.60, p_FDR_ = .045), as well as a later cluster at approximately 350–500 ms (g_z_ = −0.57, p_FDR_ = .002; Fig. 2D). Task-relevant targets, in contrast, exhibited a separate mid-latency cluster (g_z_ = 0.72, p_FDR_ = .030; Fig. 2B), and overall TRF amplitude in the attended stream was larger in magnitude across the analyzed time window.

**Figure 2.**
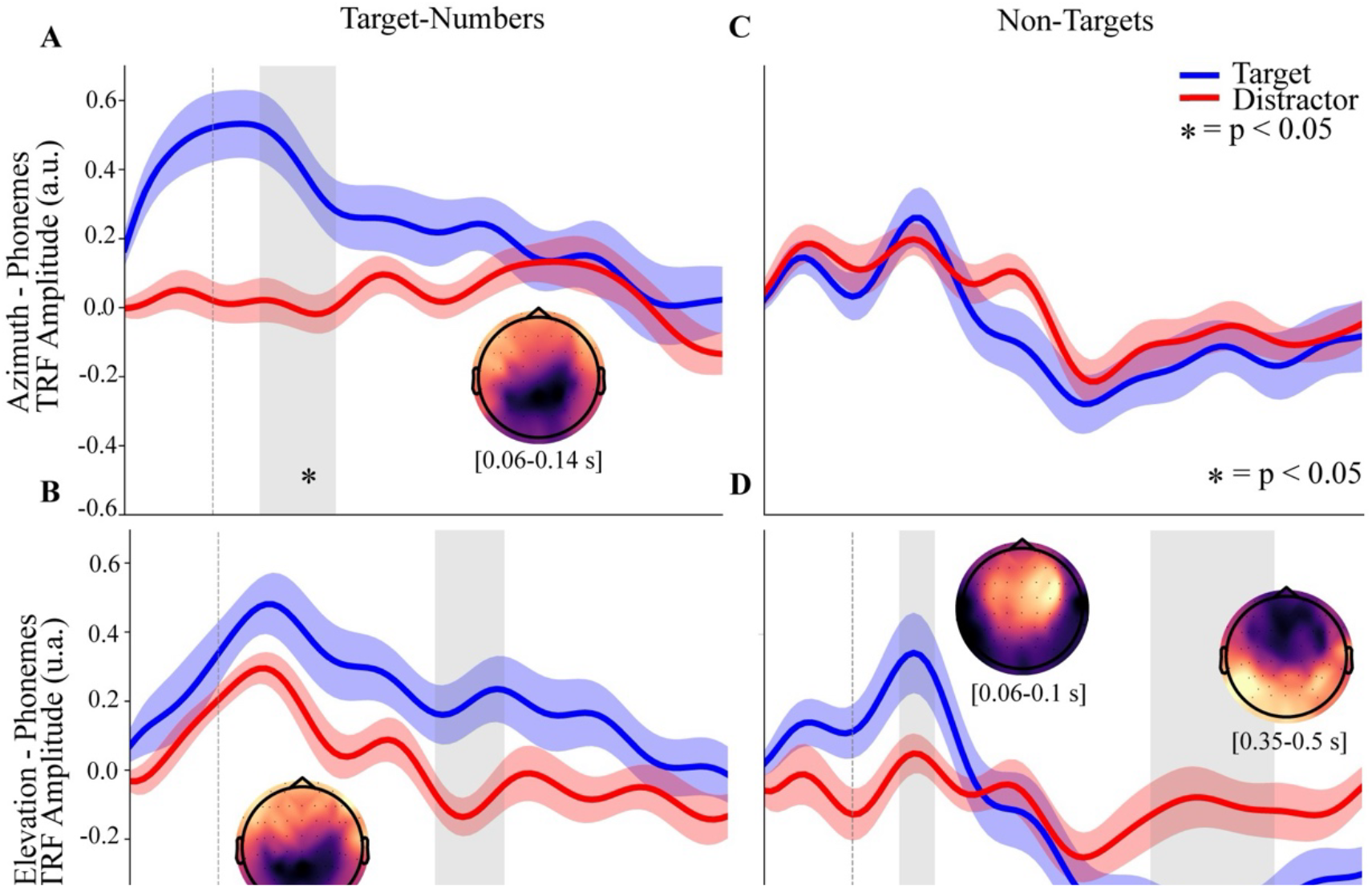
Temporal differences in TRF phoneme encoding across spatial planes and attentional streams (frontotemporal ROI). TRF amplitudes to target (A-B) and non-target (C-D) stimuli phoneme onset encoding for target (blue) and distractor (red) streams in azimuth and elevation respectively. The y-axis reflects response magnitude (a.u.; derived from standardized EEG and predictor data), and the x-axis indicates time relative to stimulus onset (–0.1 to 0.6 s). Cluster-based non-parametric permutation testing revealed a significant difference between the responses, highlighted by the shaded time window (p < .05, FDR-corrected). Insets show scalp topographies of these significant clusters (target - distractor), averaged across participants, with red indicating target > distractor and dark purple indicating distractor > target; brackets beneath each topomap indicate the corresponding latency range, rounded to two decimal places. In panel (B), no significant differences were found between target and distractor TRF responses to non-target stimuli in azimuth. The comparisons were conducted using the main, frontotemporal phoneme ROI (F3–F8, FC3–FC6, and FT7–FT8).

Because the phoneme-tracking ROI was literature-driven (Di Liberto et al., 2015: F3–F8, FC3–FC6, FT7–FT8), exploratory analyses were conducted using an alternative frontocentral ROI (FCz, F1–F6, FC1–FC6) to assess robustness. Exploratory cluster-based permutation tests using this ROI yielded qualitatively similar temporal encoding profiles (Figs. 2BD, 3), while exhibiting greater statistical sensitivity to target–distractor differences across multiple latency windows. This alternative ROI substantially overlapped with the original region but extended more centrally, suggesting that phoneme-related encoding effects may be spatially distributed and not fully captured by the predefined electrode subset. Whole-head topographic visualizations supported this interpretation, revealing broadly distributed phoneme encoding differences across frontotemporal and frontocentral scalp regions (Suppl. Figs. S1,2). Using this exploratory ROI, significant clusters were observed for target-number phoneme onsets at 96–144 ms (g_z_ = 0.48, p_FDR_ = .037), 152–200 ms (g_z_ = 0.66, p_FDR_ = .037), and 280–336 ms (g_z_ = 0.59, p_FDR_ = .037; Fig. 3A). For non-target stimuli, significant clusters were identified at 56–112 ms (g_z_ = 0.72, p_FDR_ = .035), 304–344 ms (g_z_ = −0.61, p_FDR_ = .038), and 352–496 ms (g_z_ = −0.83, p_FDR_ = .002; Fig. 3B). A similar robustness analysis was conducted for envelope encoding using a small midline cluster (FCz, Cz, CPz), selected to reflect the consistent midline distribution observed in the scalp topographies. This ROI yielded the same temporal encoding profile and significant clusters as the primary Cz analysis, confirming that envelope results were not dependent on electrode selection.

**Figure 3.**
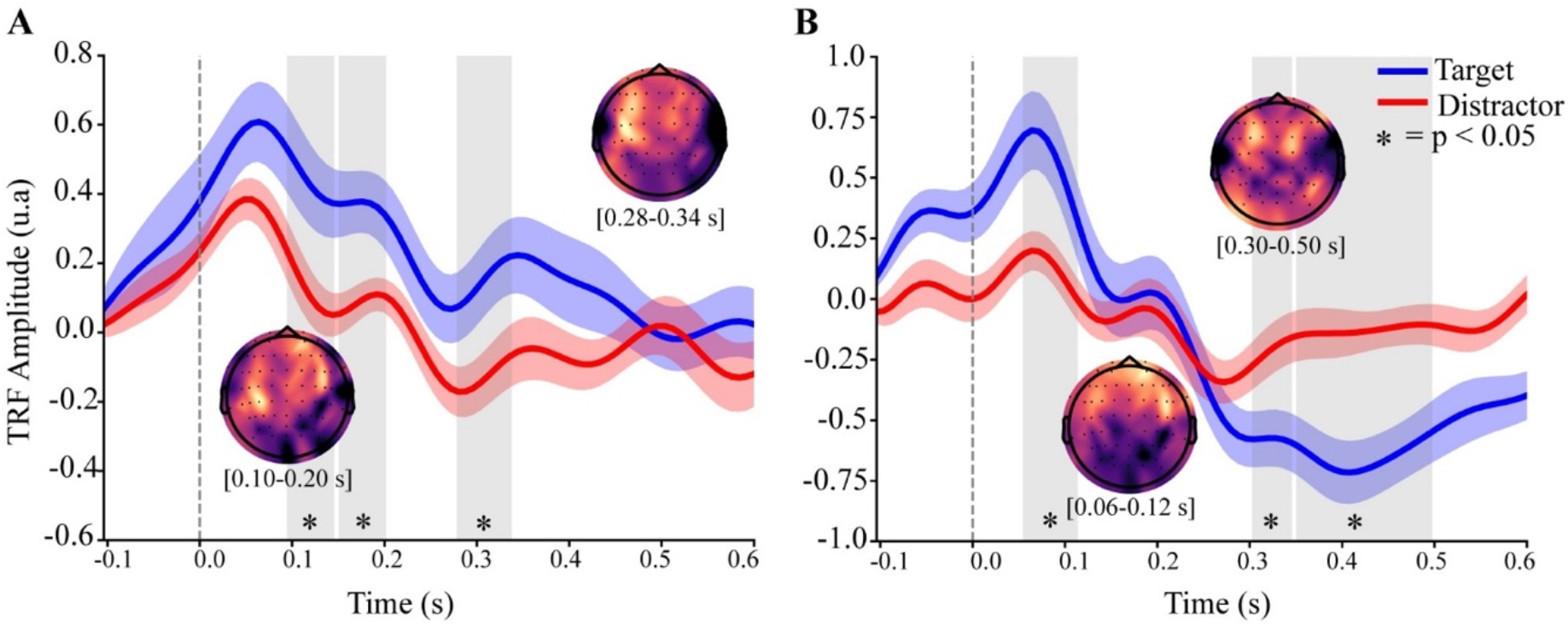
Temporal differences in TRF phoneme encoding across attentional streams (frontocentral ROI). TRF amplitudes to target (A) and non-target (B) stimuli phoneme onsets for target (blue) and distractor (red) streams in elevation. The y-axis reflects response magnitude (a.u.; derived from standardized EEG and predictor data), and the x-axis indicates time relative to stimulus onset (–0.1 to 0.6 s). Cluster-based non-parametric permutation testing revealed a significant difference between the responses, highlighted by the shaded time window (p < .05, FDR-corrected). An exploratory phoneme ROI was used, consisting of the frontocentral electrodes FCz, F1-F6 and FC1-FC6. Insets show scalp topographies of these significant clusters (target - distractor), averaged across participants, with red indicating target > distractor and dark purple indicating distractor > target; brackets beneath each topomap indicate the corresponding latency range, rounded to two decimal places.

To further confirm the robustness of these findings, cluster-based permutation tests were performed across the full 0–500 ms interval. In azimuth, envelope encoding showed significant target–distractor differences at 136–200 ms (g_z_ = 1.38, p_FDR_ = .001) and 240–312 ms (g_z_ = −1.39, p_FDR_ = .001), replicating the temporal pattern observed in window-based analyses. In elevation, phoneme encoding showed a significant early cluster at 0–112 ms (g_z_ = 0.93, p_FDR_ = .025), confirming that the strongest attentional effects remain significant under conservative whole-epoch correction.

Across-plane comparisons of the TRF target– distractor difference waves of envelope amplitudes revealed distinct temporal signatures of auditory spatial processing. Cluster-based permutation tests comparing azimuth and elevation conditions showed significant differences for non-target stimuli (Fig. 4A). In azimuth, significant across-plane differences were observed at 128–144 ms (g_z_ = 0.93, p_FDR_ = .039) and 152–184 ms (g_z_ = 1.04, p_FDR_ = .011), whereas elevation showed a distinct increase at 256–280 ms (g_z_ = −0.65, p_FDR_ = .043). No reliable across-plane differences emerged for target-number stimuli. Furthermore, using the primary frontotemporal ROI, phoneme encoding did not show significant across-plane differences (Fig. 3B). However, exploratory analyses using a frontocentral ROI revealed significant across-plane clusters at 56–144 ms (g_z_ = −0.72, p_FDR_ = .009) and 360–480 ms (g_z_ = 0.61, p_FDR_ = .025), suggesting that phoneme encoding differences may be spatially distributed and partially outside the predefined ROI.

**Figure 4.**
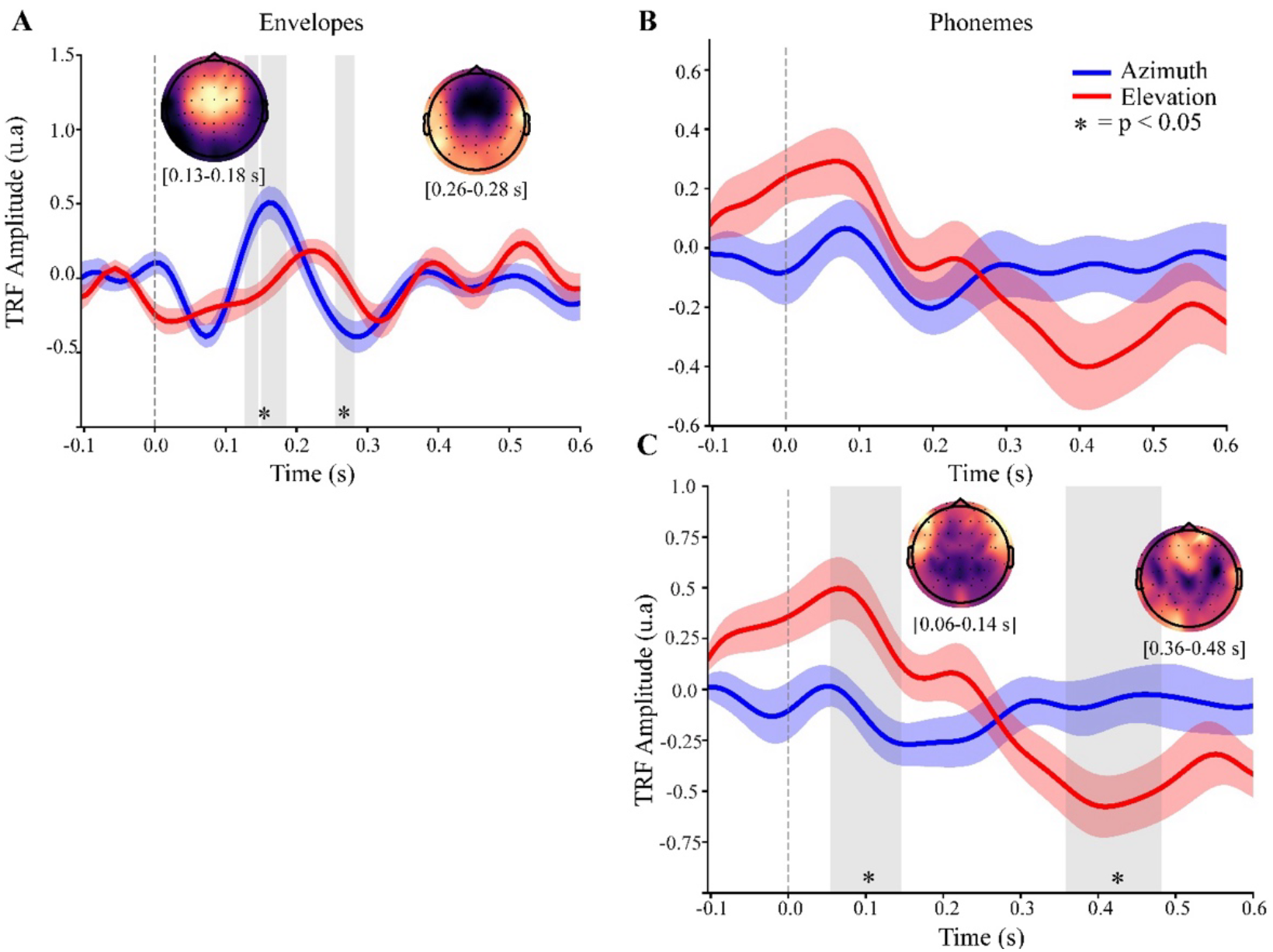
Comparison of difference waves of TRF amplitudes for envelope and phoneme onset of non-target stimuli across spatial planes. Target–distractor stream difference waves of TRF amplitudes are shown for azimuth (blue) and elevation (red), illustrating temporal differences in attentional selectivity across spatial planes. (A) Envelope predictor difference waves in the envelope ROI (Cz). (B) Phoneme predictor difference waves of the main frontotemporal ROI (F3-F8, FC3-FC6 and FT7-FT8), and (C) phoneme predictor difference waves of the exploratory frontocentral phoneme ROI (FCz, F1-F6 and FC1-FC6). Shaded gray regions indicate time windows with significant encoding differences between azimuth and elevation (p < .05, FDR-corrected). Insets show scalp topographies of these significant clusters (target - distractor), averaged across participants, with red indicating target > distractor and dark purple indicating distractor > target; brackets beneath each topomap indicate the corresponding latency range, rounded to two decimal places

There were clear differences in neural–behavioral coupling across azimuth and elevation. Linear mixed-effects models revealed significant relationships between neural selectivity (Δr) and behavioral performance in the azimuth plane for both stimulus features. For envelopes, higher Δr values predicted lower accuracy (β = −0.40, SE = 0.15, z = −2.32, p_FDR_ = .040), whereas phoneme Δr showed the opposite pattern, with stronger phonemic selectivity predicting better performance (β = 0.46, SE = 0.15, z = 3.08, p_FDR_ = .008). In both cases, the Δr × Direction interaction was nonsignificant (all p_FDR_ > .60), indicating that the neural–behavioral relationship was stable across left and right configurations. In elevation, Δr effects did not reach significance after correction (envelopes: β = −0.30, p_FDR_ = .293; phonemes: β = 0.22, p_FDR_ = .469), and directional interactions were nonsignificant (all p_FDR_ = .951). Across-plane models likewise revealed no significant Plane × Δr interactions (envelopes: β = −0.15, p_FDR_ = .389; phonemes: β = 0.32, p_FDR_ = .086; interaction terms p_FDR_ = .979). Figure 5 illustrates the relationship between neural coupling strength (Δr) and behavioral performance across predictors and spatial planes.

**Figure 5.**
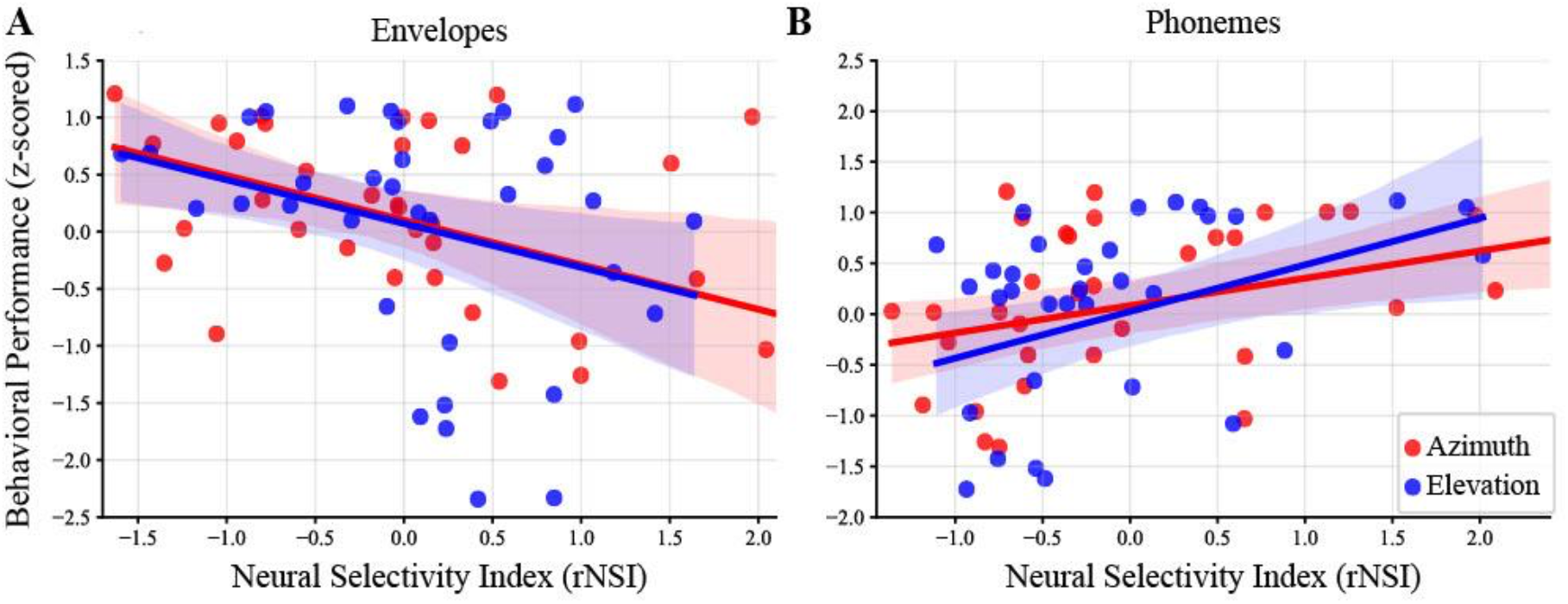
Correlation between performance accuracy and neural coupling strength (Δr) of predictors. Scatter plots showing the relationship between participants’ performance accuracy (y-axis) and the neural coupling strength Δr (x-axis), defined as the difference in model prediction accuracy (r) between target and distractor streams, for envelope (A) and phoneme onsets (B) predictors. Separate regression fits are displayed for azimuth (red) and elevation (blue), with shaded areas showing the standard error of the mean (SEM). Mixed-effects analyses revealed a significant negative relationship between performance and envelope Δr in the azimuth condition (p = .040, FDR-corrected). For the phoneme models, a positive relationship was observed, again reaching significance in the azimuth only (p = .008, FDR-corrected). In both cases the elevation followed the same trends without reaching significance (p > .05, FDR-corrected).

## DISCUSSION

Although accuracy differed across planes, performance was uniformly high overall, confirming that participants were able to perform the task reliably under both spatial configurations. According to classic spatial hearing research, azimuth trials benefitted from highly precise ITD and ILD cues, yielding near-ceiling performance even at the smallest feasible speaker separation. This aligns with early work showing that listeners can segregate sources separated by as little as 1° horizontally (Mills et al., 1958) and with later findings demonstrating that horizontal localization is supported by robust, low-variance binaural cues. Hit-rates in the elevation condition were moderately lower and more variable despite the larger speaker separation, but participants were still able to complete the task easily, with a mean hit rate of around 90%. This pattern is consistent with well-established vertical-plane limitations: minimum audible angle thresholds are substantially larger in elevation (Perrott & Saberi, 1990), spectral cues underlying vertical localization are highly dependent on individual anatomical filtering and show large variability across listeners (Giguère et al., 2011), and segregation in the vertical midline becomes significantly harder when interaural cues are minimized (Middlebrooks & Onsan, 2012). Complementary work further shows that monaural spectral cues alone, as in median-plane configurations, yield poorer localization compared to conditions where ITD/ILD information remains available (Butler & Humanski, 1992). Despite these expected differences, participants maintained high accuracy in elevation, suggesting that neural differences between planes are unlikely to reflect task disengagement or inability to perform the task, and instead likely relate to differences in spatial cue processing. It is therefore improbable that behavioral performance alone accounts for the observed differences in TRF encoding between spatial planes, and we interpret the results as the extent to which the representations of binaural and spectral cues support stream segregation.

### Early Plane-Dependent Dissociations in Envelope Encoding

Comparing the azimuth and elevation planes revealed that the mechanisms supporting selective stream tracking depend strongly on spatial configuration. In azimuth, the TRF envelope responses showed clear and temporally structured target–distractor stream differences, emerging first in early and then again at mid-latencies. We interpret the early window as reflecting stronger distractor encoding, while the later window reflects stronger target encoding. This temporal pattern closely resembles findings by Kong and colleagues (2014), who reported enhanced early P1 responses for unattended speech followed by stronger N1 responses for attended streams, reflecting differential attentional modulation across early and mid-latency processing stages. Although TRF weights do not map directly onto canonical ERP components, the correspondence in timing suggests that similar early and mid-latency attentional mechanisms shape the neural separation of competing streams in azimuth. Importantly, the dominance of envelope-driven responses in azimuth is consistent with the notion that precise binaural cues enable rapid, low-level segregation. Such early mechanisms align with bottom-up stream formation processes described by Bregman (1990) and with evidence that grouping based on primitive features such as spatial location occurs at early processing stages and can operate preattentively (Bronkhorst, 2015). Physiological studies further support this framework, showing that auditory cortical neurons represent competing spatial streams using partially distinct neural populations, enabling neural-level stream segregation (Middlebrooks & Bremen, 2013). In this context, attention may act on representations that are separated early in the auditory cortex. This helps explain why target-distractor separation emerged quickly and reliably in azimuth. Envelope encoding provides an efficient, low-level cue for tracking the target stream, enabling the system to rapidly enhance attended sounds and attenuate competing inputs within very early sensory processing stages (Woldorff et al., 1993). These results were further supported by direct across-plane comparisons of the envelope target– distractor streams amplitude differences. Azimuth showed significantly stronger target–distractor separation at early-to-mid latencies (∼150–185 ms), whereas elevation exhibited stronger separation at a later latency (∼255–280 ms). This temporal crossover was evident for non-target stimuli, while responses to target-number events did not differ across planes, presumably due to their high salience in the task context. Together, these findings indicate that envelope-driven attentional modulation follows distinct temporal profiles depending on spatial configuration.

### Similar Phoneme Encoding Across Azimuth and Elevation

In elevation, TRF responses to phoneme onsets in target-number and non-target stimuli differed between spatial streams. These differences spanned early latencies and extended beyond 0.1 s. Because the initial phoneme onset of each word was excluded from the TRF model, these early clusters are unlikely to reflect simple word onset responses. Moreover, the cluster at ∼0.25-0.35 s mirrored the late effect observed in the envelope TRF comparisons, suggesting shared later-stage processing independent of stimulus feature. These effects may reflect attention-dependent stimulus evaluation or higher-order cognitive processes, occurring approximately 200–350 ms post-stimulus (Folstein & Van Petten, 2008; Crowley & Colrain, 2004). Because button presses were explicitly modeled in the TRF, this late component is unlikely to reflect motor preparation. In contrast, the phoneme TRF responses in the azimuth condition differed between target and distractor streams only for target-number stimuli and only within an early-latency window. This pattern suggests that phoneme-level encoding in azimuth primarily reflects task-dependent attentional enhancement of behaviorally relevant stimuli. This may occur because robust binaural spatial cues already support effective stream tracking at earlier processing stages, potentially reducing the need for phoneme-level differentiation.

### Different Neural–Behavioral Coupling in Azimuth and Elevation

In the azimuth condition, the difference in TRF amplitudes between streams of both envelopes and phoneme onsets was significantly associated with behavioral performance. Greater envelope encoding differences were associated with reduced accuracy, despite overall performance being near-ceiling. Although counterintuitive, this pattern aligns with MEG findings showing that speech envelope tracking follows an inverted-U relationship with intelligibility, peaking for moderately degraded speech rather than clear speech, suggesting that increased tracking reflects compensatory attentional recruitment and listening effort rather than more efficient neural encoding (Hauswald et al., 2020). Therefore, listeners who performed best may have achieved accurate target selection with relatively efficient neural filtering, without requiring large differences in model-predicted encoding between streams. In contrast, greater phoneme encoding differences were associated with better performance, aligned with evidence that stronger neural encoding of linguistic content reflects more effective speech comprehension and predicts behavioral intelligibility (Broderick et al., 2018). Overall, these opposing trends suggest two distinct mechanisms: individuals who experience the task as more difficult may rely more heavily on low-level acoustic segregation, reflected in increased envelope encoding difference, whereas stronger phoneme-level selectivity may reflect more efficient linguistic processing, and therefore better overall speech intelligibility and performance. This interpretation, however, remains speculative and should be confirmed in tasks producing greater variability in behavioral performance, where neural–behavior relationships can be assessed across a broader range of attentional success.

With envelope encoding showing clear and temporally robust target–distractor separation in azimuth, spanning multiple latency windows, the findings align with previous work, demonstrating that selective attention enhances neural encoding of acoustic features, including speech envelopes and temporal modulations (Lakatos et al., 2005; Xiang et al., 2010; Mesgarani & Chang, 2012). Conversely, envelope-driven selectivity in elevation was weaker and limited to a later latency window, and phoneme-level encoding within the elevation plane exhibited more widespread attentional modulation across time. This may suggest a greater relative contribution of phonemic representations when acoustic spatial cues are less informative.

The lack of significant phoneme encoding differences between spatial planes in the primary analysis may partly reflect the conservative, literature-based ROI selection. Statistical comparisons of TRF phoneme amplitudes showed increased sensitivity when using the exploratory frontocentral ROI, suggesting that phoneme-level attentional modulation in elevation may not be optimally captured by the predefined frontotemporal ROI. Consistent with this interpretation, neural–behavioral coupling analyses using the primary phoneme ROI revealed similar directional relationships between phoneme selectivity and performance across planes, but these effects did not reach statistical significance in elevation. This pattern is unlikely to reflect greater inter-individual variability in phoneme encoding, as phoneme TRF prediction accuracy exhibited lower inter-subject variability than envelope models across both spatial planes. Instead, the absence of statistical significance in elevation likely reflects reduced sensitivity to smaller phoneme-related effects. Several methodological factors may have contributed to this reduced sensitivity. First, although the sample size is comparable to prior TRF studies, smaller effects, particularly those involving higher-level linguistic features, may require larger samples to detect reliably, especially in experimental designs that reduce low-level acoustic entrainment and remove the availability of voice-related cues supporting selective stream tracking. Second, phoneme predictors were implemented as onset impulse trains and therefore do not capture richer linguistic representations, which may have limited sensitivity to higher-order encoding differences. Future studies using larger samples and richer linguistic predictors may therefore help to further clarify how attentional modulation of phonemic representations supports stream tracking when spatial cues are less reliable.

Together, these findings suggest that speech features may be differentially modulated across spatial planes, with envelope encoding playing a more dominant role in stream tracking along azimuth, whereas phoneme encoding may contribute more when binaural spatial cues provide less reliable support, reflecting adaptive neural mechanisms that flexibly prioritize acoustic and phonemic representations depending on spatial cue reliability.

## Availability Statement

The datasets generated and analyzed for this study are not publicly available due to participant privacy and data protection regulations but are available from the corresponding author upon reasonable request.

## Author Contributions

VKK: Conceptualization, Methodology, Investigation, Project Administration, Resources (participant recruitment), Formal Analysis, Visualization, Writing – Original Draft.

MS: Conceptualization, Resources (laboratory instrumentation), Supervision, Validation, Writing – Review & Editing, Funding Acquisition (via existing lab support).

Both authors contributed to manuscript revision, approved the submitted version, and agreed to be accountable for the work.

---

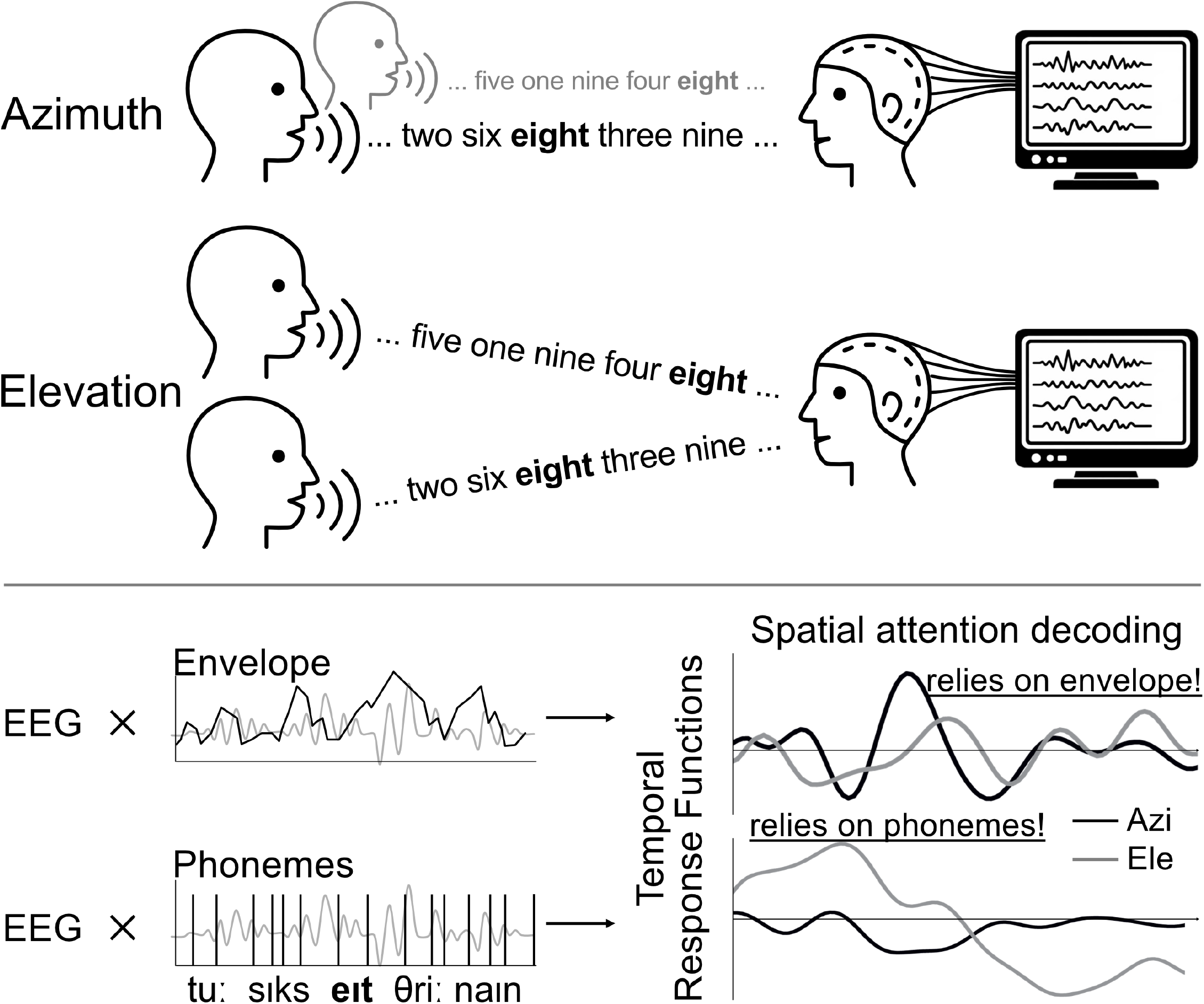

## References

1. E. Macaluso, C.D. Frith, J. Driver, Directing Attention to Locations and to Sensory Modalities: Multiple Levels of Selective Processing revealed with PET, Cerebral Cortex, Volume 12, Issue 4, April 2002, Pages 357–368, 10.1093/cercor/12.4.357

2. Shomstein, S., & Yantis, S. (2004). Control of attention shifts between vision and audition in human cortex. The Journal of neuroscience 24(47), 10702–10706

3. Fu, D., Weber, C., Yang, G., Kerzel, M., Nan, W., Barros, P., Wu, H., Liu, X., & Wermter, S. (2020). What Can Computational Models Learn From Human Selective Attention? A Review From an Audiovisual Unimodal and Crossmodal Perspective. Frontiers in integrative neuroscience, 14, 10

4. Daniel, P. M., & Whitteridge, D. (1961). The representation of the visual field on the cerebral cortex in monkeys. The Journal of Physiology, 159, 203–221. 10.1113/jphysiol.1961.sp006803

5. Dow, B. M., Snyder, A. Z., Vautin, R. G., & Bauer, R. (1981). Magnification factor and receptive field size in foveal striate cortex of the monkey. Experimental brain research, 44(2), 213–228. 10.1007/BF00237343

6. Cherry, E. C. (1953). Some experiments on the recognition of speech, with one and with two ears. J. Acoust. Soc. Am. 25, 975–979.

7. Bregman, A. S. (1990). Auditory Scene Analysis: The Perceptual Organization of Sound. Cambridge, MA: The MIT Press. 10.7551/mitpress/1486.001.0001

8. Sussman, E., Ritter, W. and Vaughan, H.G. (1999) ‘An investigation of the auditory streaming effect using event-related brain potentials’, Psychophysiology, 36(1), pp. 22–34. doi:10.1017/S0048577299971056.

9. Sussman, E.S., Bregman, A.S., Wang, W.J. et al. Attentional modulation of electrophysiological activity in auditory cortex for unattended sounds within multistream auditory environments. Cognitive, Affective, & Behavioral Neuroscience 5, 93–110 (2005). 10.3758/CABN.5.1.93

10. Best, V., Ozmeral, E. J., Kopco, N., & Shinn-Cunningham, B. G. (2008). Object continuity enhances selective auditory attention. Proceedings of the National Academy of Sciences of the United States of America, 105(35), 13174–13178. 10.1073/pnas.0803718105

11. O’Sullivan, J. A., Power, A. J., Mesgarani, N., Rajaram, S., Foxe, J. J., Shinn-Cunningham, B. G., Slaney, M., Shamma, S. A., & Lalor, E. C. (2015). Attentional Selection in a Cocktail Party Environment Can Be Decoded from Single-Trial EEG. Cerebral cortex (New York, N.Y. : 1991), 25(7), 1697–1706. 10.1093/cercor/bht355

12. Ding, N., & Simon, J. Z. (2012). Emergence of neural encoding of auditory objects while listening to competing speakers. Proceedings of the National Academy of Sciences of the United States of America, 109(29), 11854–11859. 10.1073/pnas.1205381109

13. Zion Golumbic, E. M., Ding, N., Bickel, S., Lakatos, P., Schevon, C. A., McKhann, G. M., Goodman, R. R., Emerson, R., Mehta, A. D., Simon, J. Z., Poeppel, D., & Schroeder, C. E. (2013). Mechanisms underlying selective neuronal tracking of attended speech at a “cocktail party”. Neuron, 77(5), 980–991. 10.1016/j.neuron.2012.12.037

14. Broderick, M. P., Anderson, A. J., Di Liberto, G. M., Crosse, M. J., & Lalor, E. C. (2018). Electrophysiological correlates of semantic dissimilarity reflect the comprehension of natural, narrative speech. Current Biology, 28(5), 803–809.e3. 10.1016/j.cub.2018.01.080

15. Power, A. J., Foxe, J. J., Forde, E.-J., Reilly, R. B., & Lalor, E. C. (2012). At what time is the cocktail party? A late locus of selective attention to natural speech. European Journal of Neuroscience, 35(9), 1497–1503. 10.1111/j.1460-9568.2012.08060.x

16. Kaufman, M., & Zion Golumbic, E. (2023). Listening to two speakers: Capacity and tradeoffs in neural speech tracking during Selective and Distributed Attention. NeuroImage, 270, 119984. 10.1016/j.neuroimage.2023.119984

17. Middlebrooks, J. C., & Green, D. M. (1991). Sound localization by human listeners. Annual Review of Psychology, 42, 135–159. 10.1146/annurev.ps.42.020191.001031

18. Blauert, J. (1997). Spatial hearing: the psychophysics of human sound localization. 10.7551/mitpress/6391.001.0001

19. Deng, Y., Choi, I., Shinn-Cunningham, B., & Baumgartner, R. (2019). Impoverished auditory cues limit engagement of brain networks controlling spatial selective attention. NeuroImage, 202, 116151. 10.1016/j.neuroimage.2019.116151

20. Mills, A. W. (1958). On the minimum audible angle. Journal of the Acoustical Society of America, 30, 237–246. 10.1121/1.1909553

21. Masterton B, Diamond IT. 1967. Medial superior olive and sound localization. Science 155(770):1696–7

22. Yin, T. C. & Chan, J. C. Interaural time sensitivity in medial superior olive of cat. J. Neurophysiol. 64, 465–488 (1990)

23. Yao JD, Bremen P, Middlebrooks JC. Emergence of Spatial Stream Segregation in the Ascending Auditory Pathway. J Neurosci. 2015 Dec 9;35(49):16199–212.

24. Batteau DW. The role of the pinna in human localization. Proc R Soc Lond B Biol Sci. 1967 Aug 15;168(1011):158–80.

25. Wightman FL, Kistler DJ (1989) Headphone simulation of free-field listening: II. Psychophysical validation. J Acoust Soc Am 85:868–878.

26. Xu L, Furukawa S, Middlebrooks JC. Sensitivity to sound-source elevation in nontonotopic auditory cortex. J Neurophysiol. 1998 Aug;80(2):882–94.

27. Crosse, M. J., Di Liberto, G. M., Bednar, A., & Lalor, E. C. (2016). The multivariate temporal response function (mTRF) toolbox: A MATLAB toolbox for relating neural signals to continuous stimuli. Frontiers in Human Neuroscience, 10, Article 604. 10.3389/fnhum.2016.00604

28. Schönwiesner & Bialas (2021). s(ound)lab: An easy to learn Python package for designing and running psychoacoustic experiments. Journal of Open Source Software 6.62, p. 3284

29. Gramfort, A., Luessi, M., Larson, E., Engemann, D. A., Strohmeier, D., Brodbeck, C., Goj, R., Jas, M., Brooks, T., Parkkonen, L., & Hämäläinen, M. (2013). MEG and EEG data analysis with MNE-Python. Frontiers in neuroscience, 7, 267. 10.3389/fnins.2013.00267

30. Bialas et al. (2023). mTRFpy: A Python package for temporal response function analysis. Journal of Open Source Software, 8(89), 5657. 10.21105/joss.05657

31. McAuliffe, M., Socolof, M., Mihuc, S., Wagner, M., Sonderegger, M. (2017) Montreal Forced Aligner: Trainable Text-Speech Alignment Using Kaldi. Proc. Interspeech 2017, 498–502, doi: 10.21437/Interspeech.2017-1386

32. Aiken, S. J., & Picton, T. W. (2008). Envelope and spectral frequency-following responses to vowel sounds. Hearing Research, 245(1–2), 35–47. 10.1016/j.heares.2008.08.004

33. Lalor, E. C., & Foxe, J. J. (2010). Neural responses to uninterrupted natural speech can be extracted with precise temporal resolution. European Journal of Neuroscience, 31(1), 189–193.

34. Di Liberto, G. M., O’Sullivan, J. A., & Lalor, E. C. (2015). Low-Frequency Cortical Entrainment to Speech Reflects Phoneme-Level Processing. Current biology : CB, 25(19), 2457–2465. 10.1016/j.cub.2015.08.030

35. Benjamini, Y., & Hochberg, Y. (1995). Controlling the false discovery rate: A practical andpowerful approach to multiple testing. Journal of the Royal Statistical Society B, 57,289–300

36. Perrott, D. R., & Saberi, K. (1990). Minimum audible angle thresholds for sources varying in both elevation and azimuth. The Journal of the Acoustical Society of America, 87(4), 1728–1731. 10.1121/1.399421

37. Giguere, C., Lavallee, R., Plourde, J., & Vaillancourt, V. (2011). Vertical sound localization in left, median and right lateral planes. Canadian Acoustics, 39(4), 3–12.

38. Middlebrooks, J. C., & Onsan, Z. A. (2012). Stream segregation with high spatial acuity. The Journal of the Acoustical Society of America, 132(6), 3896–3911. 10.1121/1.4764879

39. Butler, R.A., Humanski, R.A. Localization of sound in the vertical plane with and without high-frequency spectral cues. Perception & Psychophysics 51, 182–186 (1992). 10.3758/BF03212242

40. Kong, Y. Y., Mullangi, A., & Ding, N. (2014). Differential modulation of auditory responses to attended and unattended speech in different listening conditions. Hearing research, 316, 73–81. 10.1016/j.heares.2014.07.009

41. Bronkhorst, A.W. The cocktail-party problem revisited: early processing and selection of multi-talker speech. Atten Percept Psychophys 77, 1465–1487 (2015). 10.3758/s13414-015-0882-9

42. Middlebrooks, J. C., & Bremen, P. (2013). Spatial stream segregation by auditory cortical neurons. The Journal of neuroscience, 33(27), 10986–11001. 10.1523/JNEUROSCI.1065-13.2013

43. Woldorff, M. G., Gallen, C. C., Hampson, S. A., Hillyard, S. A., Pantev, C., Sobel, D., & Bloom, F. E. (1993). Modulation of early sensory processing in human auditory cortex during auditory selective attention. Proceedings of the National Academy of Sciences of the United States of America, 90(18), 8722–8726. 10.1073/pnas.90.18.8722

44. Folstein, J. R., & Van Petten, C. (2008). Influence of cognitive control and mismatch on the N2 component of the ERP: a review. Psychophysiology, 45(1), 152–170. 10.1111/j.1469-8986.2007.00602.x

45. Crowley, K. E., & Colrain, I. M. (2004). A review of the evidence for P2 being an independent component process: age, sleep and modality. Clinical neurophysiology, 115(4), 732–744. 10.1016/j.clinph.2003.11.021

46. Hauswald, A., Keitel, A., Chen, Y. P., Rösch, S., & Weisz, N. (2022). Degradation levels of continuous speech affect neural speech tracking and alpha power differently. The European journal of neuroscience, 55(11-12), 3288–3302. 10.1111/ejn.14912

47. Lakatos, P., Shah, A. S., Knuth, K. H., Ulbert, I., Karmos, G., & Schroeder, C. E. (2005). An oscillatory hierarchy controlling neuronal excitability and stimulus processing in the auditory cortex. Journal of neurophysiology, 94(3), 1904–1911. 10.1152/jn.00263.2005

48. Xiang, J., Simon, J., & Elhilali, M. (2010). Competing streams at the cocktail party: exploring the mechanisms of attention and temporal integration. The Journal of neuroscience, 30(36), 12084–12093.

49. Mesgarani, N., & Chang, E. F. (2012). Selective cortical representation of attended speaker in multi-talker speech perception. Nature, 485(7397), 233–236. 10.1038/nature11020

