## Supplemental Figures for "Speech Stream Tracking in 2D: Attention Differentially Enhances Acoustic and Phonemic Encoding Across Spatial Planes"

### Supplementary Material

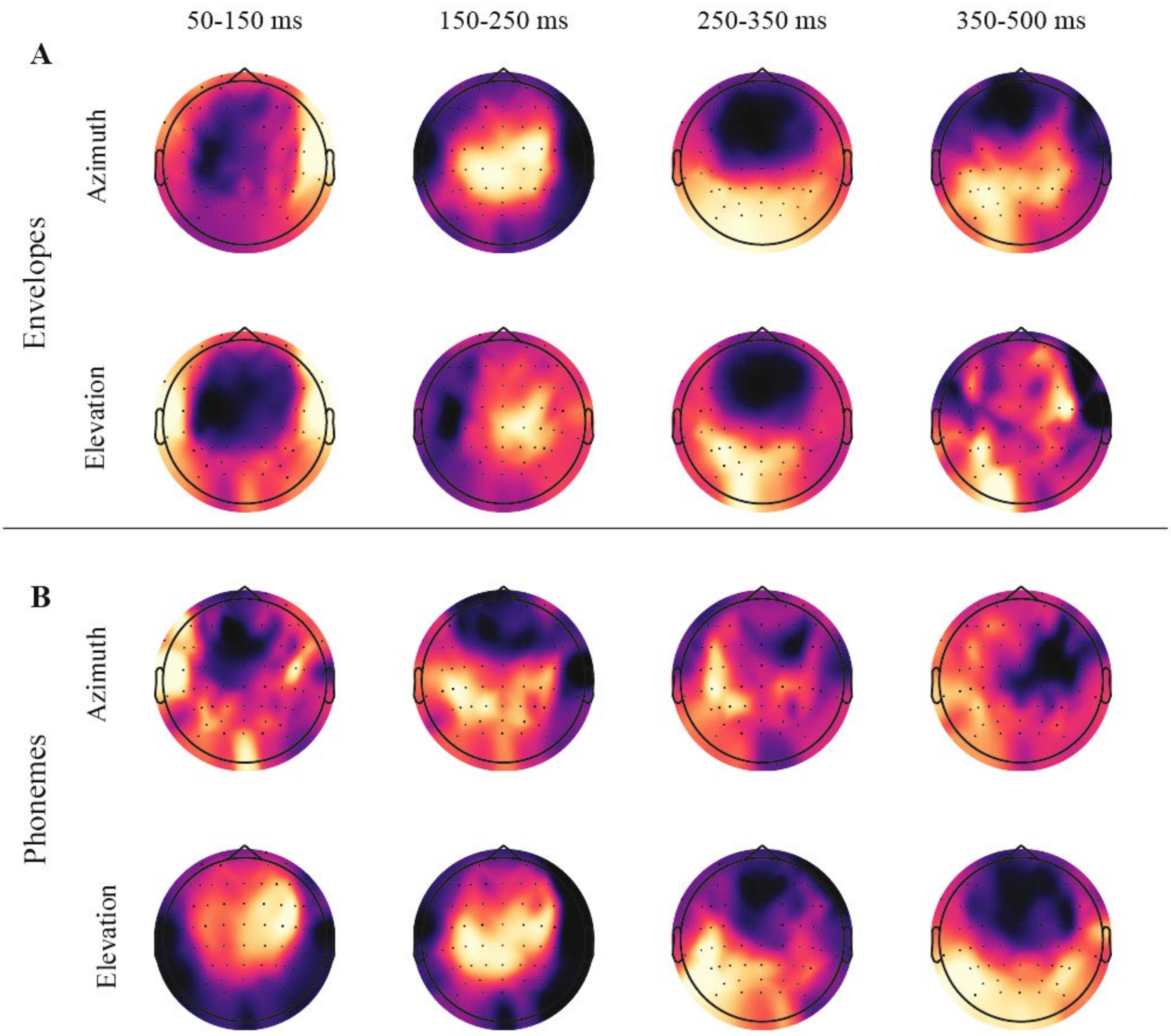


**Figure S1: Topographic distribution of Δamplitude between predicted TRF responses for target and distractor streams (target–distractor), computed across subjects for target-number stimuli. (A)** Envelope and **(B)** phoneme onsets predictors. Each panel shows topographic maps for azimuth and elevation conditions, across four time-windows: 50–150 ms, 150–250 ms, 250–350 ms, and 350–500 ms. Δamplitude reflects the z-scored difference in predicted TRF response amplitude between target and distractor streams. All EEG channels are shown. Regions of interest (ROIs) were defined based on prior literature: the phoneme ROI comprised fronto-temporal electrodes (F3–F8, FC3–FC6, FT7–FT8; Di Liberto et al., 2015), and the envelope ROI was defined at Cz. Yellow indicates greater target > distractor, whereas blue distractor > target.

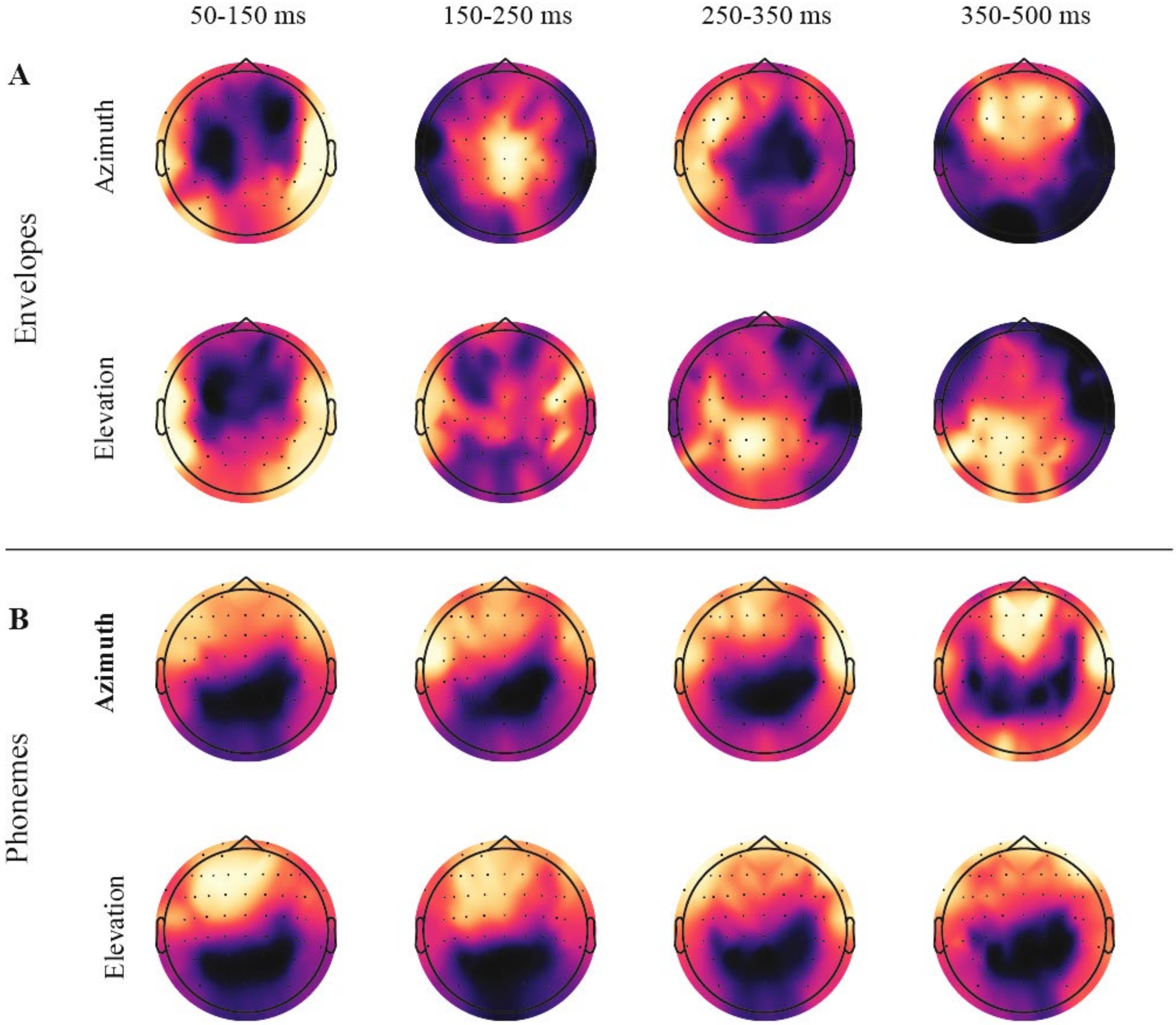


**Figure S2: Topographic distribution of Δamplitude between predicted TRF responses for target and distractor streams (target–distractor), computed across subjects for non-target number stimuli. (A)** Envelope and **(B)** phoneme onsets predictors. Each panel shows topographic maps for azimuth and elevation conditions, across four time-windows: 50–150 ms, 150–250 ms, 250–350 ms, and 350–500 ms. Δamplitude reflects the z-scored difference in predicted TRF response amplitude between target and distractor streams. All EEG channels are shown. Regions of interest (ROIs) were defined based on prior literature: the phoneme ROI comprised fronto-temporal electrodes (F3–F8, FC3–FC6, FT7–FT8; Di Liberto et al., 2015), and the envelope ROI was defined at Cz. Yellow indicates greater target > distractor, whereas blue distractor > target.

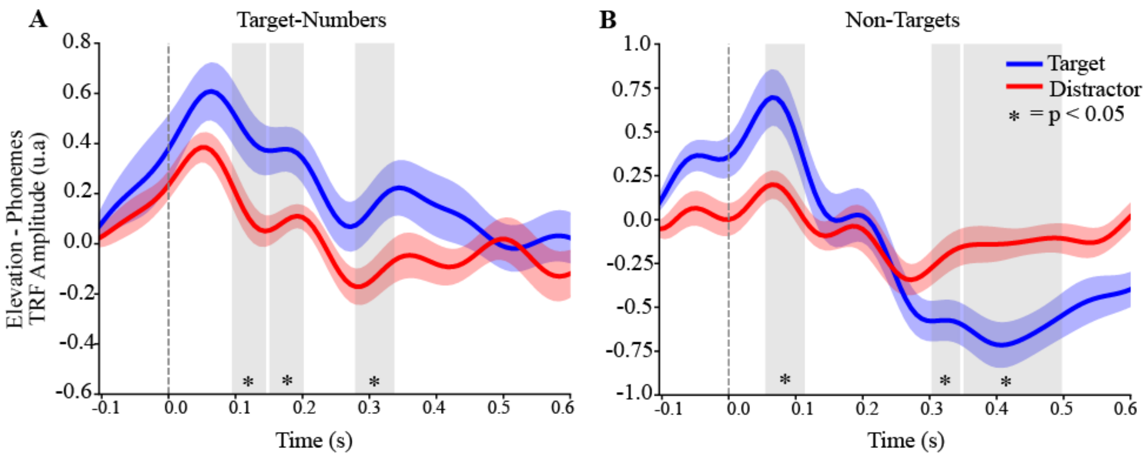


**Figure S3: Phoneme encoding of target-number and non-target stimuli in elevation across frontocentral electrodes.** TRF predicted encoding to target **(A)** and non-target **(B)** stimuli from phoneme onsets for target (blue) and distractor (red) streams in elevation. The y-axis reflects response magnitude (a.u.; derived from standardized EEG and predictor data), and the x-axis indicates time relative to stimulus onset (–0.1 to 0.6 s). Cluster-based non-parametric permutation testing revealed a significant difference between the responses, highlighted by the shaded time window ($p < 0.05$). In panel **(A)**, three significant clusters were observed for target-number phoneme onsets (96-144 ms, $g_z$=0.481, $p_{FDR}$=0.037; 152-200 ms, $g_z$=0.661, $p_{FDR}$=0.037; 280-336 ms, $g_z$=0.593, $p_{FDR}$=0.037. In panel **(B)**, three significant clusters were observed for non-targets phoneme onsets (56-112 ms, $g_z$=0.721, $p_{FDR}$=0.035; 304-344 ms, $g_z$=-0.610, $p_{FDR}$=0.038; 352-496 ms; $g_z$=-834, $p_{FDR}$ = 0.002). The exploratory phonemes ROI used consisted of the frontocentral electrodes FCz, F1-F6 and FC1-FC6, with substantial overlap with the main, literature-driven ROI (Di Liberto et al., 2015: F3-F8, FC3-FC6 and FT7-FT8).

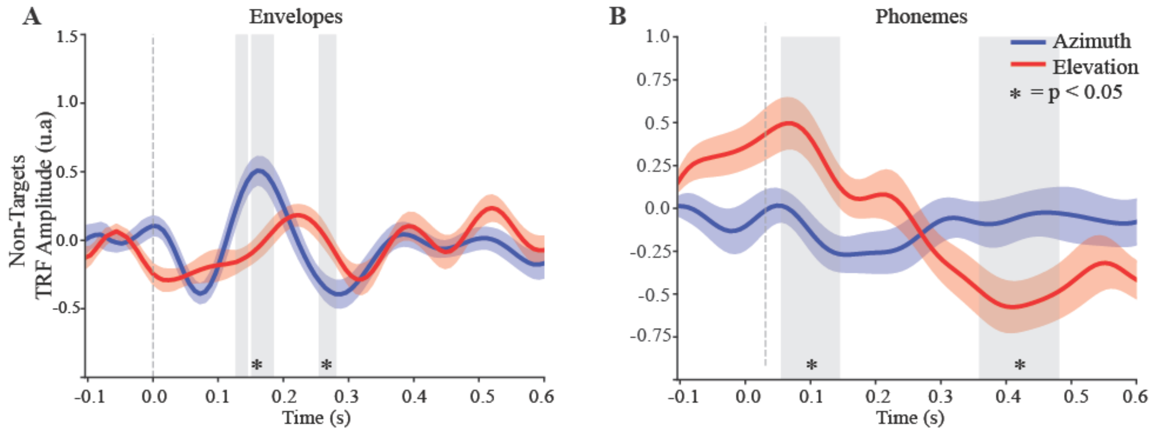


**Figure S4: Predictor encoding of non-target stimuli across planes across frontocentral electrodes.** Difference waves of TRF predicted responses (target-distractor) to envelopes **(A)** and phoneme onsets **(B)** in azimuth (blue) and elevation (red). The y-axis reflects response magnitude (a.u.; derived from standardized EEG and predictor data), and the x-axis indicates time relative to stimulus onset (–0.1 to 0.6 s). Shaded regions indicate significant differences between planes, identified using cluster-based non-parametric permutation testing ($p < 0.05$, FDR-corrected). In panel **(A)**, three significant clusters were observed for envelope predictors (128-144 ms, $g_z$=0.932, $p_{FDR}$=0.039; 152-184 ms, $g_z$=1.040, $p_{FDR}$=0.011; 256-280 ms, $g_z$=-0.646, $p_{FDR}$=0.043), computed at electrode Cz, consistent with the envelope ROI used in the main analysis. In panel **(B)**, two significant clusters were observed for phoneme onsets (56-144 ms, $g_z$=-0.723, $p_{FDR}$=0.009; 360-480 ms, $g_z$=0.607, $p_{FDR}$=0.025). The exploratory phonemes ROI used consisted of the frontocentral electrodes FCz, F1-F6 and FC1-FC6, with substantial overlap with the main, literature-driven ROI (Di Liberto et al., 2015: F3-F8, FC3-FC6 and FT7-FT8).
